# Loss of mitochondrial SIRT4 shortens lifespan and leads to a decline in physical activity

**DOI:** 10.1101/248831

**Authors:** Sweta Parik, Sandipan Tewary, Champakali Ayyub, Ullas Kolthur-Seetharam

**Author notes:** Corresponding author: Ullas Kolthur-Seetharam (*). **Authors contact information:** Sweta Parik, Sandipan Tewary, Champakali Ayyub.

## Abstract

Mitochondrial mechanisms and pathways have recently emerged as critical determinants of organismal aging. While nuclear sirtuins have been shown to regulate aging, whether mitochondrial sirtuins do so is still unclear. Here, we report that mitochondrial *dSirt4* mediates organismal survival. We establish that absence of *dSirt4* leads to reduced lifespan independent of dietary inputs. Further by assaying locomotion, a key correlate of aging, we demonstrate that *dSirt4* null flies are severely physically impaired with a significant reduction in locomotion. In summary, we report for the first time that mitochondrial *dSirt4* is a key determinant of longevity and its loss leads to early aging.

---

Genetic mutations that impair mitochondrial function have been associated with a shortened lifespan (Sun *et al*. 2016). However, the importance of a regulatory factor such as a mitochondrial metabolic sensor in regulating aging is still not known.

Sirtuins are evolutionarily conserved NAD+ dependent enzymes, which impinge on cellular functions by deacylating several proteins (Chang & Guarente 2014). SIR2/SIRT1 and SIRT6 are essential determinants of organismal survival and lifespan (Rogina & Helfand 2004; Banerjee *et al*. 2012; Kanfi *et al*. 2012; Satoh *et al*. 2013). We have earlier demonstrated that *dSir2/Sirt1* positively regulates lifespan in a diet dependent manner, using control and *dSir2/Sirt1* perturbed flies, which were otherwise genetically identical (Banerjee *et al*. 2012).

Mitochondrial sirtuins have emerged as key regulators of several mitochondrial functions, which contribute to healthy aging and organismal survival (He *et al*. 2012). It should be noted that among mitochondrial sirtuins, SIRT4 is one of the least characterized across species. Others and we have demonstrated that SIRT4 is a negative regulator of catabolic processes and is crucial for maintaining cellular ATP (Nasrin *et al*. 2010; Ho *et al*. 2013). Although, SIRT4 null mice display enhanced fatty acid oxidation (Nasrin *et al*. 2010; Laurent *et al*. 2013), there are no reports on long-term consequences of SIRT4 absence on lifespan.

In this context, we sought to determine the role of SIRT4 in aging using *Drosophila melanogaster* as a model system. Phylogenetic analyses indicated that dSIRT4 is the fly orthologue of human SIRT4 (Figure 1A). We also generated transgenic flies expressing Myc-tagged dSIRT4 and found it to be exclusively localized to the mitochondria like its mammalian counterpart (Figure 1B).

We set out to determine the effects of both loss-and gain-of-function of *dSirt4* on lifespan using a deletion mutant (Figure 1C) and UAS-dSirt4-Myc expressing flies (Figure 1D). Transgenic UAS-*dSirt4*-Myc flies showed more than 40 fold increase in *dSirt4* expression (Figure 1D). Since supra-physiological expression of proteins may lead to non-physiological phenotypes, we did not pursue assaying for lifespan in these flies. In order to negate potentially confounding background genetic variations, we generated backcrossed *dSirt4* mutants (hereafter denoted as *dSirt4*^-/-*bck*^).

**Figure 1:**
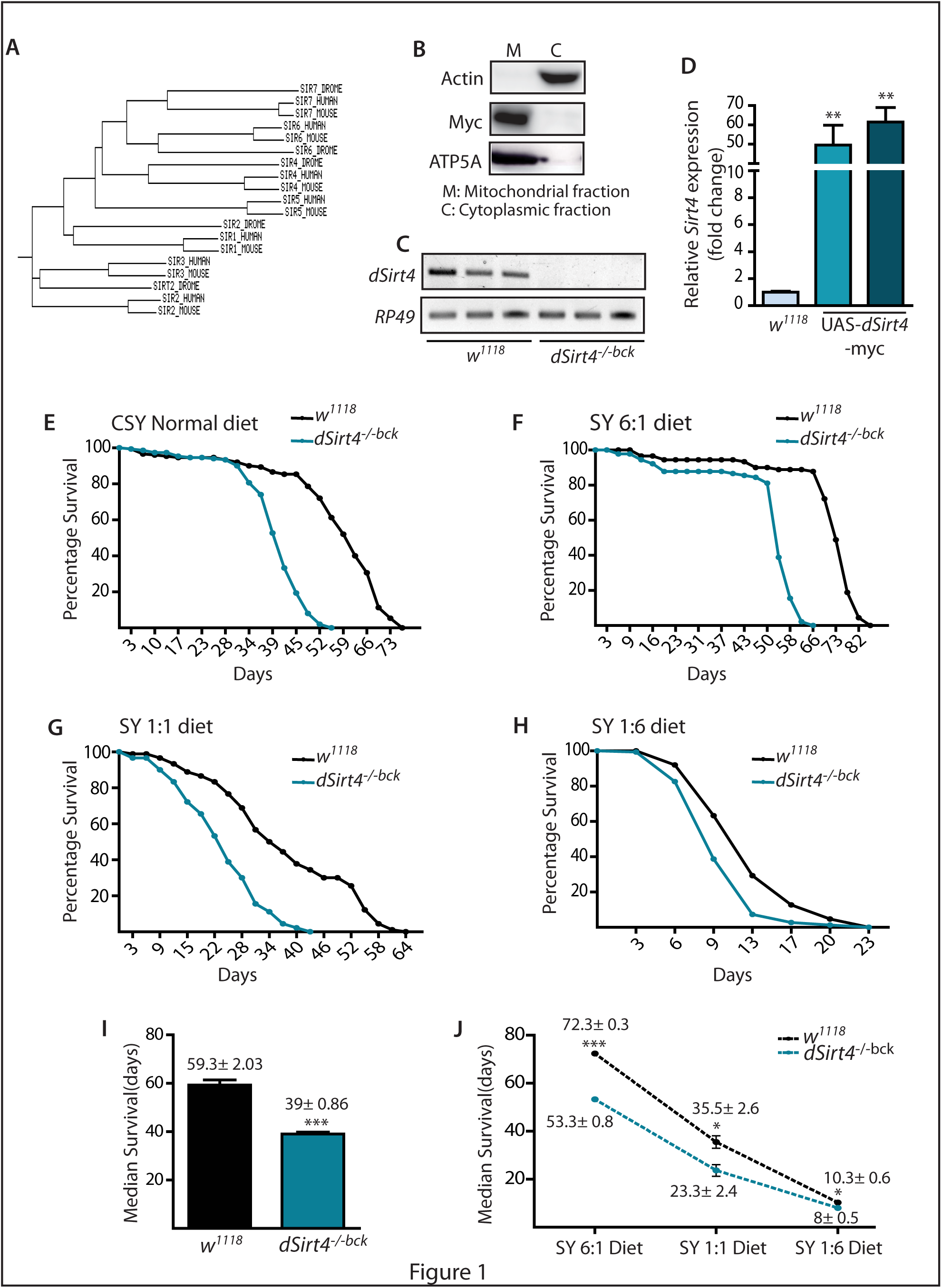
*dSirt4*^-/-*bck*^ flies have reduced lifespan across dietary conditions-. (A) Phylogeny of sirtuins (SIRT1-7) in the indicated species (B) Mitochondrial localisation of dSIRT4-MYC in transgenic flies (C) *dSirt4* mRNA expression in control and *dSirt4*^-/-*bck*^ flies (D) *dSirt4* mRNA expression in control and *UAS-dSirt4* flies. (E-J) Lifespan of control and *dSirt4*^-/-*bck*^ flies reared under (E) Standard cornmeal diet (N=2, n=160) (F) S:Y 6:1 diet (N=2, n=160) (G) S:Y 1:1 diet (N=2, n=160) and (H) S:Y 1:6 diet (N=3, n=220). Median lifespan of control and *dSirt4*^-/-*bck*^ under (I) standard diet and (J) S:Y media.

On standard cornmeal diet, *dSirt4*^-/-*bck*^ flies had a significantly shorter lifespan when compared to control flies (Figures 1E and 1I). Relative ratios of sugar and yeast in diet rather than their concentration determine organismal physiology and longevity (Lee *et al*. 2008; Zhu *et al*. 2014). On assaying for survival under varying dietary regimes (sugar:yeast or S:Y diets), we found that *dSirt4*^-/-*bck*^ flies had significantly shortened lifespan on all the diets (Figure 1F-H). It is important to note that increasing yeast concentrations led to early death in control flies as reported earlier (Lee *et al*. 2008). However, absence of *dSirt4* led to a further reduction in lifespan across diets (Figure 1J). Moreover, these results emphasize that a background genetic mutation would unlikely contribute to the reduced longevity phenotype across all dietary regimes.

The progressive loss of muscle mass and hence strength, with age, termed “sarcopenia” is often deemed responsible for weakness and impaired locomotion in aged individuals (Cruz-Jentoft *et al*. 2010). In *Drosophila*, while most tissues exhibit age-induced damage, several studies have shown that thoracic muscles, required for walking/climbing and flight display the most severe defects (Demontis *et al*. 2013).

To check if reduced lifespan was associated with accelerated aging, we assessed the physical activity of both control and *dSirt4*^-/-*bck*^ flies by performing a climbing assay (Figure 2A). We then quantified climbing efficiency in terms of relative positions in the vial at different time points and percentage of flies that cross a threshold distance at a particular time point (Supplementary information). As shown in Figures 2B and 2C, we saw that control flies showed an age-dependent decrease in locomotion as previously reported. In 8-10 days old flies, there was a minor lag in the climbing activity of *dSirt4*^-/-*bck*^ flies at 15 seconds with a higher percentage of flies occupying region 2 and lesser percentage of flies occupying region 4, as compared to controls (Figure 2B). This difference was not observed when the activity was quantified at later time points (Figure 2B).

**Figure 2:**
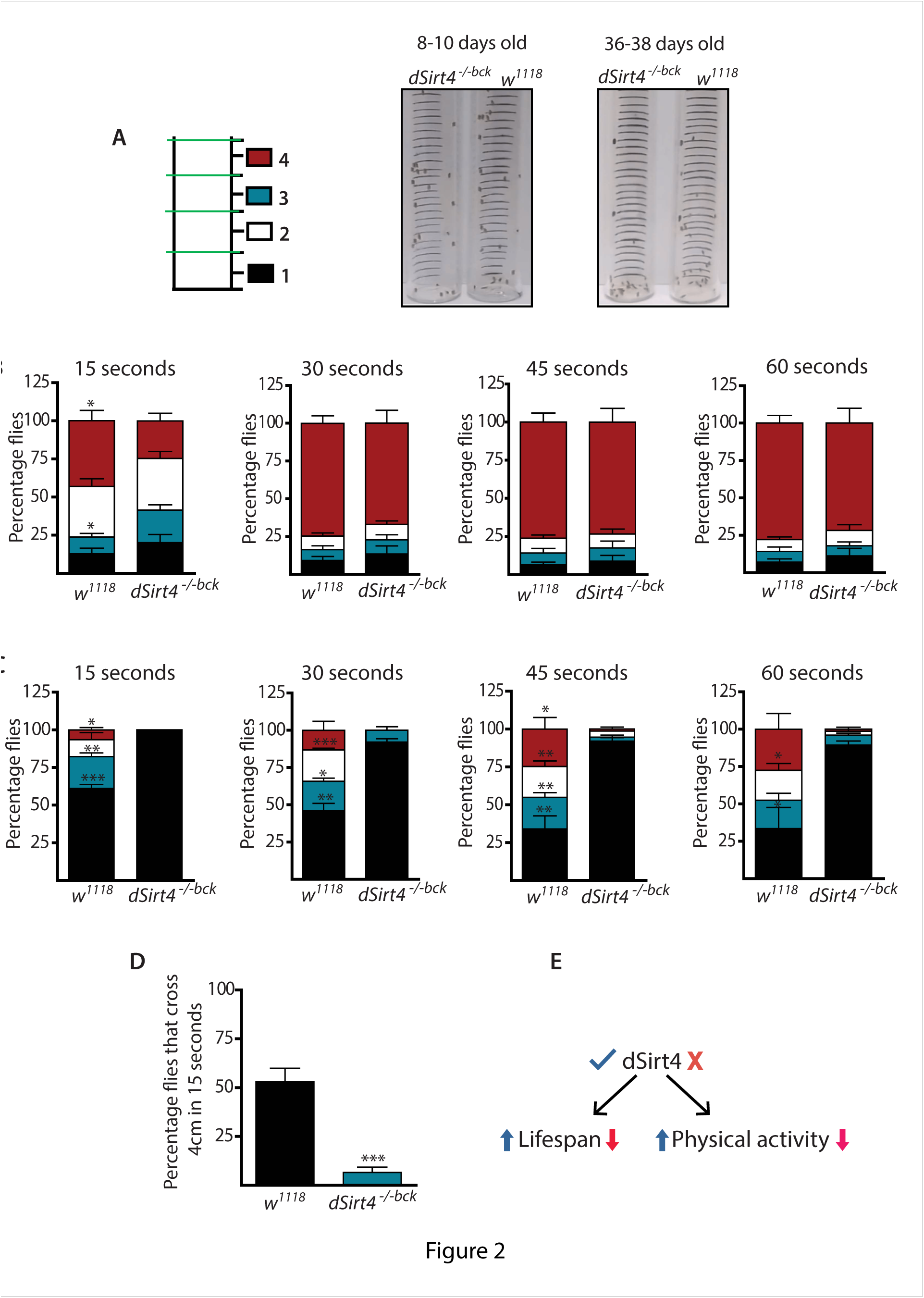
Aged *dSirt4*^-/-*bck*^ flies have reduced physical activity-. (A) Schematic and representative images from video for negative geotactic climbing assay and quantification. (B-C) Climbing activity in control and *dSirt4*^-/-*bck*^ flies, which were (B) 8-10 days old (N=4, n=285) and (C) 36-38 days old (N=3, n=210). (D) Percentage of 36-38 days old control and *dSirt4*^-/-*bck*^ flies that cross a distance of 4cm in 15 seconds. (E) *dSirt4* regulates lifespan and physical activity.

Importantly, we found a striking decrease in climbing activity in *dSirt4*^-/-*bck*^ flies when compared to age matched 36-38 days old control flies (Figure 2C). The drastic decline was made apparent by the fact that most mutant flies occupied the base of the vial (region 1) and a significantly smaller percentage was seen at the top (regions 3 & 4). *dSirt4*^-/-*bck*^ flies were found to be severely impaired in their ability to cross a threshold distance (Figure 2E).

Taken together, our results demonstrate that *dSirt4^-/-bck^* flies have reduced climbing ability with a severe reduction in locomotion, which is enhanced in aged cohorts. It also clearly indicates that absence of *dSirt4* leads to a loss of physical activity akin to early aging.

In conclusion, our novel findings illustrate that *dSirt4*, a mitochondrial sirtuin, is essential for organismal survival across diets and that its loss leads to accelerated aging. In the future, it will be exciting to assess the molecular and genetic mechanisms that mediate these *dSirt4* dependent effects on aging, and in a tissue specific manner.

## Acknowledgements

We acknowledge Namrata Shukla, Arpan Parichha and Kushal Banerjee for technical help.

## Author Contributions

S.P., S.T. and C.A. carried out experiments; U.K.S. conceptualized the project with inputs from S.P. and C.A.; Manuscript was written by S.P. and U.K.S. U.K.S. obtained the funds and supervised the project.

